# Electrophysiological properties and synaptic activity of the mouse hippocampal CA1 neurons during postnatal development

**DOI:** 10.64898/2026.06.15.732242

**Authors:** Igor Nagula, Emilija Kavalnyte, Kornelija Vitkute, Daiva Dabkeviciene, Urte Neniskyte, Aidas Alaburda

## Abstract

Early postnatal development is a critical period for hippocampal circuit maturation. While postnatal hippocampal development has been mostly studied in rats, less is known about the developmental trajectory of electrophysiological properties in mice, despite the wide use of these animal models for molecular and genetic studies of nervous system. In this study, we investigated the postnatal maturation of hippocampal CA1 pyramidal neurons in male and female wild-type mice. Whole-cell patch-clamp recordings were performed in acute hippocampal slices to assess passive and active membrane properties as well as spontaneous excitatory synaptic activity. We found that maturation of neuronal firing properties was associated with faster responses to stimulation, higher-amplitude and shorter-duration action potentials, and more precise control of neuronal firing. Simultaneously, synaptic activity changed across development, with decreased sEPSC inter-event intervals and stable event amplitudes, suggesting enhanced functional connectivity without major changes in synaptic strength. Sex-dependent differences in electrophysiological properties were observed primarily during the first postnatal week, indicating that sex influences the early trajectory of neuronal maturation. Together, our findings provide a comprehensive electrophysiological baseline for mouse hippocampal CA1 pyramidal neurons during postnatal development.

## Introduction

The hippocampus, a key part of the limbic system, plays a crucial role in long-term and spatial memory, navigation, and learning processes (Bangasser and Shors, 2007; Burgess et al., 2002). The postnatal development of the hippocampus is crucial, as it involves neuronal transformation, the formation of new neural circuits, synaptic maturation, plasticity, and myelination (Yang et al., 2024; Ábrahám et al., 2010; Reznikov, 1991; Pokorný and Yamamoto, 1981). Moreover, abnormalities in hippocampal development, such as aberrant pruning, are linked to various neurodevelopmental disorders, including autism spectrum disorders, epilepsy and schizophrenia (Neniskyte and Gross, 2017). The first three weeks of postnatal development of hippocampal neurons are characterized by morphological and electrophysiological changes in rats and mice due to the expression and incorporation of new ion channels and receptors (Sánchez-Aguilera et al., 2020; Moody and Bosma, 2005). These changes can be linked to emerging exploratory behaviour in rodents after eye opening between postnatal days P12 and P15 (Hoy and Niell, 2015; Wills et al., 2010; Lu and Constantine-Paton, 2004). Moreover, maturation of intrinsic electrophysiological properties of CA1 neurons coincides with the period of synaptic proliferation and refinement and may be associated with the peak in synaptogenesis of CA1 neurons, which occurs between two and four weeks after birth (Yang et al., 2024; Paolicelli et al., 2011; Wills et al., 2010). Electrophysiological studies on brain development in rats have demonstrated significant changes in membrane properties of neurons, including input resistance, membrane time constant, action potential threshold, width, amplitude, and upstroke/downstroke velocities in the cerebral cortex and hippocampal CA1 pyramidal neurons during postnatal development (Brahimi et al., 2023; Sánchez-Aguilera et al., 2020; Giglio and Storm, 2014; Sengupta et al., 2013; Spigelman et al., 1992; McCormick and Prince, 1987). These findings indicate that both active and passive intrinsic electrophysiological properties undergo substantial changes during development in rats. However, changes in intrinsic electrophysiological properties during the development in mice have been studied primarily in cortical regions, such as the anterior cingulate cortex, medial prefrontal cortex, and primary visual cortex (Ciganok-Hückels et al., 2023; Kroon et al., 2019; Pan et al., 2016). Consequently, it remains unclear how the electrophysiological properties, information processing capabilities and synaptic activity of mouse hippocampal CA1 pyramidal neurons change during early postnatal development. Moreover, morphological and immunohistochemical studies show sex differences in cell proliferation during postnatal brain development in the hippocampal CA1, CA3, and dentate gyrus regions (Bowers et al., 2010; Zhang et al., 2008). Previous studies have also demonstrated that sex hormones influence the density of dendritic spines in hippocampal CA1 pyramidal neurons, with these changes occurring during maturation in male rats and across the estrous cycle in female rats (Gould et al., 1990; Woolley et al., 1990; Meyer et al., 1978). These findings underscore the need to consider sex differences in electrophysiological studies. In this study, we examined the passive and active electrophysiological properties, firing properties and the synaptic activity of mouse hippocampal CA1 pyramidal neurons during critical postnatal development in both sexes using whole-cell patch clamp electrophysiology.

This study enhances the understanding of postnatal hippocampal development and provides a baseline for electrophysiological studies of postnatal hippocampal maturation in mice.

## Materials and Methods

### Animals

C57BL/6J wild-type mice were obtained from local Vilnius University Life Sciences Center animal facility colony. Animals were housed in groups and maintained on a 12- hour light/dark cycle under consistent conditions of temperature (21.5 ± 1 °C) and humidity (55 ± 8%). Access to water and food was provided *ad libitum*. Animal experiments were performed at Vilnius University with the approval by the State Food and Veterinary Service of the Republic of Lithuania (No. G2-91). Efforts were made to minimize animal suffering and to reduce the number of laboratory animals used, in accordance with the European Communities Council Directive of September 20, 2010 (2010/63/EU).

### Experimental design

Electrophysiological data were obtained from mouse hippocampal slices. In total, 51 animals, both males and females, were used in experiments. The animals were categorized into groups of different postnatal age stages: P4-P6 (P5), P8-P9 (P8), P13- P16 (P15), and P18-P23 (P21). Furthermore, each group was further subdivided by sex into male and female cohorts. 262 neurons were analysed, from 1 to 15 neurons from each animal.

### Acute brain slice preparation

Acute brain slice preparation was adapted from a N-methyl-d-glucamine (NMDG) protective recovery method, described previously (Ting et al., 2018). Animals were anesthetized by inhalation of isoflurane and immediately decapitated. The brain was dissected out in a cold and carbonated NMDG-HEPES artificial cerebrospinal fluid (aCSF) solution containing: 92 mM NMDG, 2.5 mM KCl, 1.25 mM NaH_2_PO_4_, 30 mM NaHCO_3_, 20 mM HEPES, 25 mM Glucose, 2 mM Thiourea, 5 mM Na-ascorbate, 3 mM Na-pyruvate, 0.5 mM CaCl_2_·2H_2_O, 10 mM MgSO_4_·7H_2_O, pH 7.3–7.4 (95% O_2_/5% CO_2_ and 300–310 mOsmol/kg). The rostral and caudal parts of the brain were cut with a scalpel. The brain was dried with filter paper and mounted on an angled 12° holder of a vibratome (*5100mz Vibratome, Campden Instruments Ltd*), with the ventral side positioned down and the rostral part facing up the angle. Hippocampal slices (300 μm- thick) were cut with the vibratome in a cold and oxygenated NMDG-HEPES aCSF solution and transferred to a chamber filled with pre-warmed (32–34 °C) and continuously carbonated NMDG-HEPES aCSF. Within the 10 minutes of transferring the acute brain slices, 2 M NaCl solution was gradually introduced into the NMDG HEPES aCSF, following the Na^+^ spike-in procedure (Ting et al., 2018). After the Na^+^ spike-in procedure was complete, the slices were maintained for at least 1 hour in a continuously oxygenated, room-temperature HEPES holding aCSF solution containing: 92 mM NaCl, 2.5 mM KCl, 1.25 mM NaH_2_PO_4_, 30 mM NaHCO_3_, 20 mM HEPES, 25 mM glucose, 2 mM thiourea, 5 mM Na-ascorbate, 3 mM Na-pyruvate, 0.5 mM CaCl_2_·2H_2_O, 4 mM MgSO_4_·7H_2_O, pH 7.3–7.4 (95% O_2_/5% CO_2_ and 300–310 mOsmol/kg).

### Whole-cell patch-clamp recordings

To record electrophysiological data, acute brain slices were transferred to a perfusion chamber and secured with a brain slice anchor. The brain slice was continuously perfused with room temperature carbonated HEPES recording aCSF (2 mL/min–5 mL/min), containing: 124 mM NaCl, 2.5 mM KCl, 1.25 mM NaH_2_PO_4_, 24 mM NaHCO_3_, 12.5 mM glucose, 5 mM HEPES, 2 mM CaCl_2_·2H_2_O, and 1 mM MgSO_4_·7H_2_O, pH 7.3–7.4 (95% O_2_/5% CO_2_ and 300–310 mOsmol/kg). Hippocampal CA1 pyramidal neurons were recorded using the patch-clamp whole-cell configuration method. Patch micropipettes (3–8 MΩ) were pulled from borosilicate glass (*1B150F-4, World Precision Instruments*) with a horizontal puller (*Puller P1000, Sutter Instruments*) and filled with freshly thawed and filtered intracellular solution, containing: 130 mM K-gluconate, 4 mM KCl, 0.3 mM EGTA, 10 mM HEPES, 10 mM Na_2_-phosphocreatine, 13.4 mM biocytin, 4 mM Mg-ATP, 0.3 mM Na_2_-GTP, pH 7.3–7.4 (290–300 mOsmol/kg). Recording location was identified using a 4× objective on a brightfield microscope. Subsequently, a 40× water-immersion objective was used to identify individual neuron bodies in the pyramidal cell layer of the hippocampus CA1b area. Whole-cell patch-clamp recordings were performed after the formation of a Gigaohm seal. Ideally, recordings were performed on cells located >30 μm deep in the slice, as there was lower risk for the neurons being physically damaged with severed dendritic processes. Only medium-sized cells with a homogeneous surface appearance and soft membrane boundaries were selected for patching. Data were recorded at a sampling rate of 20 kHz using a 16-bit analog-to-digital converter (*Digidata 1440A, Molecular Devices*) and amplifier (*MultiClamp 700B, Axon Instruments*). Recordings were displayed using *pCLAMP* software. Neurons with a resting membrane potential below a standard firing threshold (negative to -55 mV) were accepted for further analysis. Experiments were performed within 8 hours after slicing. Animal age or sex was known during recordings.

### Passive properties

Passive membrane properties were measured under current-clamp conditions. A current was injected into the cell using a step waveform function, increasing in magnitude with every step. The steps of the injected current ranged from −60 to 390 pA. Each step lasted 1 second. The resting membrane potential was measured under current-clamp conditions with no current injection. The input resistance (*R*_in_) was calculated from voltage deflection in a current-clamp configuration using the formula *R*_in_ = ΔV/ΔI. The membrane time constant (*τ*_m_) was obtained by fitting a single exponential curve to the voltage deflection.

### Active properties

Active membrane properties were obtained from the recordings in current-clamp mode, as described above. Since the values of the action potential (AP) parameters can depend on the AP frequency adaptation (Venance and Glowinski, 2003), it was chosen to measure the AP parameters at initial and steady frequencies. The initial frequency was calculated from the first interspike interval and the steady frequency was calculated as the average of the last three interspike intervals of the AP train. The initial (5–15 Hz) and steady (3–13 Hz) frequency intervals were selected according to the frequency and current relation graphs of each neuron. Initial AP parameter values were calculated from the first evoked AP of a train. Steady AP parameters were calculated as the average values of the last three APs of a train. AP threshold values were found as membrane potential values corresponding to the peak values of a third derivative of membrane potential with respect to time. AP amplitude was measured as a difference between the AP peak and the minimum membrane potential point during hyperpolarization. AP width was defined as the duration of AP at the half-maximal amplitude. AP maximal upstroke and downstroke velocities were found as peak and minimum values of the first derivative of membrane potential with respect to time. The relationship between the frequency of action potentials and the injected current (frequency-current relation, f-I) was characterized by gain. Accordingly, for each neuron, its initial gain was obtained by extracting a slope value from a linear fit of the f-I relation from the first interspike interval, while steady gain was obtained by extracting a slope value from a linear fit of the f-I relation from the last three interspike intervals. The spike-frequency adaptation ratio was defined as the ratio of the initial to the steady-state firing rate. For each neuron, the trace with an initial firing frequency closest to ∼20 Hz was selected, and the adaptation ratio was calculated from that trace.

### Synaptic activity

Spontaneous excitatory postsynaptic currents (sEPSC) were recorded by holding the membrane potential of the neuron at -70 mV for 10 minutes in a voltage-clamp configuration. Recordings were visually inspected for signal stability prior to analysis, and data segments or entire recordings exhibiting abnormal or burst-like synaptic activity were excluded. Accepted recordings were low-pass filtered at 500 Hz using a Gaussian filter and analyzed using *Clampfit* software (*Molecular Devices*). sEPSC were detected using a template-based approach, with templates created from 20–30 representative single events. Event detection was performed using the *Clampfit* template search function.

### Data analysis and statistics

Electrophysiological data were analyzed using *Clampfit* 10.7 and a custom [Python script](https://github.com/Kavalnyte-Emilija/action-potential-analysis.git) for event detection. Statistical tests were performed with *GraphPad Prism 10.1.2* (*GraphPad, San Diego, CA, USA*). Data were tested for normal distribution using the Shapiro-Wilk test. sEPSC interevent intervals and amplitudes exhibited non-normal distributions and were log-transformed prior to parametric statistical analysis. Statistical significance between group means was calculated using two-way ANOVA followed by Holm-Sidak’s multiple comparisons *post hoc* test. To compare the mean difference between two related measures, the Wilcoxon signed-rank test was used. In addition, distribution-based analyses of synaptic activity were performed using normalized histograms and cumulative probability plots to assess differences in event distributions not captured by average values. Differences between distributions were evaluated using the two-sample Kolmogorov–Smirnov (KS) test. Statistical significance was determined at *P* < 0.05. *n* represents the sample size. All values are given as mean values ± standard deviation (SD).

## Results

### The development of passive electrophysiological properties

It is known that the input resistance and time constant of other types of neurons, such as pyramidal neurons of medial prefrontal cortex in mice and cerebral cortical pyramidal neurons in rats, decrease during postnatal development, possibly because of an increase in the density of ionotropic channels (Kroon et al. 2019; McCormick and Prince 1987). Therefore, we evaluated the passive electrophysiological properties of hippocampal CA1 pyramidal neurons during postnatal development. Developmental stages are shown as postnatal days (P5, P8, P15, P21) (Table A1). Resting membrane potential was stable from P5 to P21 both in females and males. Sex-related differences appeared at P8, as male neurons showed a significantly more depolarized resting membrane potential (*P* = 0.004). (Fig. 1A; Table A1). Input resistance decreased from P5 to P21 by 2.6-fold in females (*P* < 0.001) and by 4.1-fold in males (*P* < 0.001). Sex-related differences appeared at P5, as male neurons showed a significantly higher input resistance (*P* < 0.001) (Fig. 1B; Table A1). Membrane time constant became shorter from P5 to P21 by 1.42-fold in females (*P* = 0.02) and by 1.6-fold in males (*P* < 0.001). There were no significant differences in membrane time constant between the sex groups (*P* = 0.13) (Fig. 1C; Table A1). Overall, a large decrease in input resistance and membrane time constant values suggests major changes in neuronal membrane composition during the development of the mouse hippocampus, likely reflecting ion channel incorporation and associated changes in membrane conductance.

**Fig. 1.**
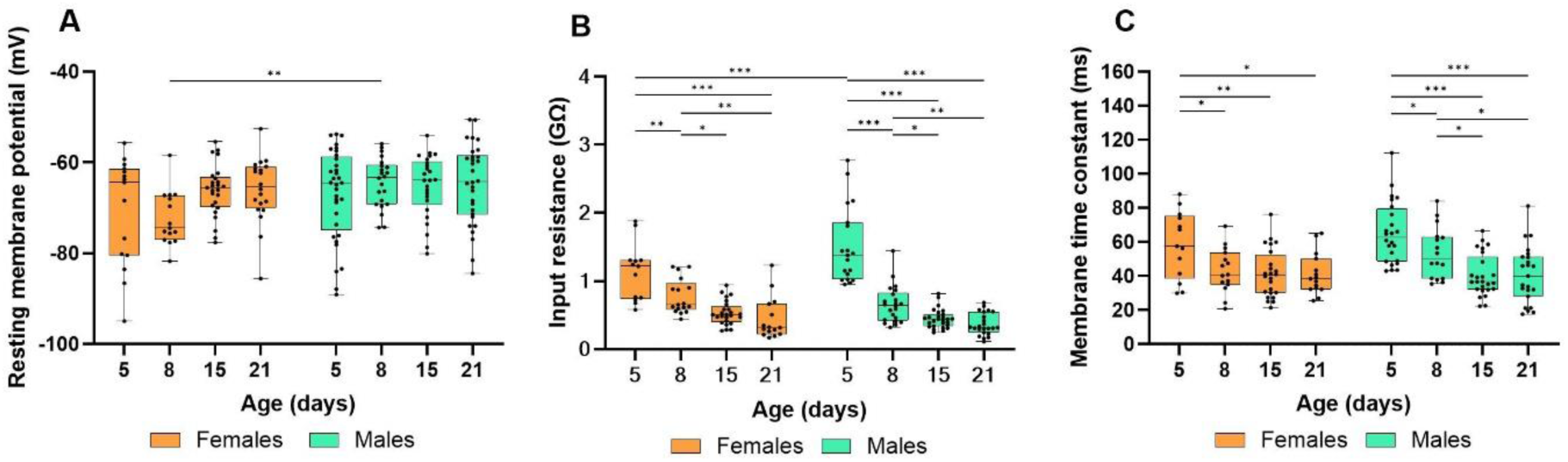
Changes in passive membrane properties during development in females and males. Developmental stages are shown as postnatal days (P5, P8, P15, P21). Asterisks indicate significant differences between groups (\**P* < 0.05, \*\**P* < 0.01, \*\*\**P* < 0.001). Orange indicates females; green indicates males. (**A**) Resting membrane potential remained stable across development, with a sex difference appearing at P8. (**B**) Input resistance and (**C**) membrane time constant decreased with age; input resistance showed a sex difference at P5.

### The development of active electrophysiological properties

The development of passive electrophysiological properties described above, together with increased expression of ion channels, may alter the active properties of neurons. Simultaneously, a variety of neurons exhibit spike frequency adaptation, characterized by a reduction in firing frequency during sustained depolarization. Previous studies have shown that this reduction in firing rate affects active membrane properties, including depolarization of the action potential (AP) threshold, decreased AP amplitude and membrane time constant, and altered spike initiation dynamics (Yi et al., 2017; Miles et al., 2005; Venance and Glowinski, 2003). In the present study, we observed spike frequency adaptation that was independent of mouse age or sex, with firing frequency decreasing by approximately 2-fold from the initial to the steady-state phase across all age groups with no significant sex-dependent differences (*P* = 0.23) (Fig. 2). Based on these observations, we evaluated active AP electrophysiological properties separately before (initial; Table A2) and after (steady; Table A3) the spike frequency adaptation.

**Fig. 2.**
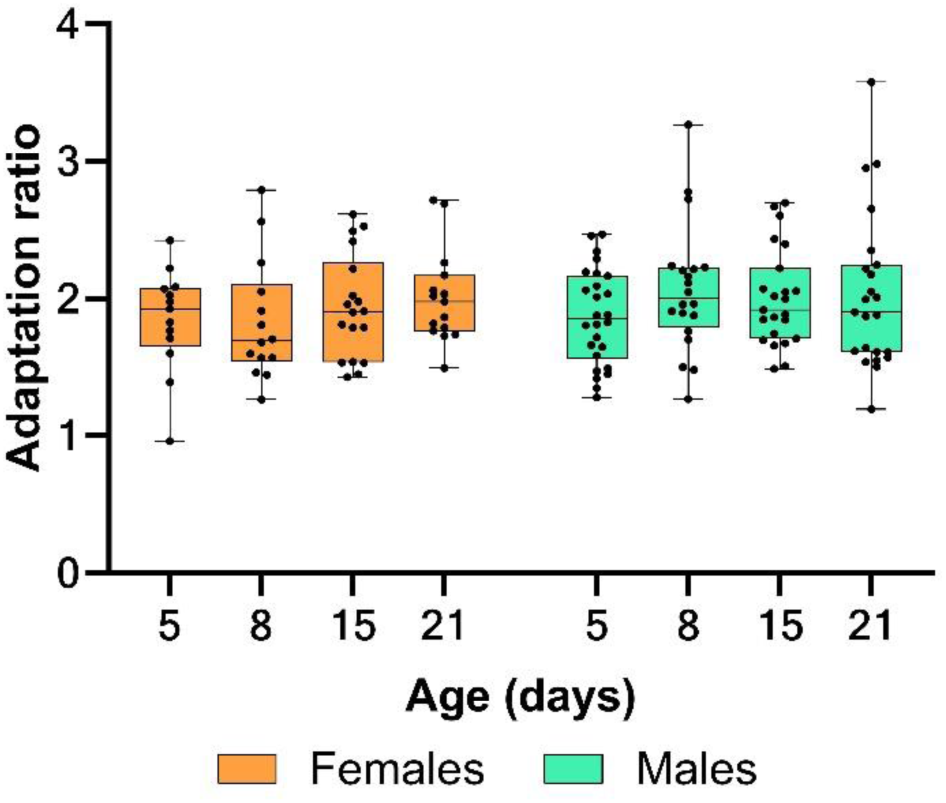
The ratio of spike frequency adaptation. Developmental stages are shown as postnatal days (P5, P8, P15, P21). Orange indicates females; green indicates males. Spike frequency decreased by approximately 2-fold after adaptation in all age groups, with no significant differences between sexes.

### Initial AP electrophysiological properties

Initial AP threshold hyperpolarised from P5 to P21 by 1.57-fold in females (*P* = 0.002) and by 2.08-fold in males (*P* < 0.001). Sex-related differences appeared at P5 (*P* = 0.02) and P8 (*P* = 0.04) as male’s neurons showed a higher initial AP threshold (Fig. 3A; Table A2). Initial AP amplitude increased from P5 to P21 by 16.2% in females (*P* = 0.001) and by 26.4% in males (*P* < 0.001). There were no significant differences in initial AP amplitude between the sex groups (*P* = 0.93) (Fig. 3B; Table A2). Initial AP width decreased from P5 to P21 by 1.61-fold in females (*P* < 0.001) and by 1.85-fold in males (*P* < 0.001). There were no differences in initial AP width between the sex groups (*P* = 0.11) (Fig. 3C; Table A2). Initial maximal AP upstroke velocity increased from P5 to P21 by 1.79-fold in females (*P* < 0.001) and by 2.05-fold in males (*P* < 0.001). There were no sex differences in initial maximal AP upstroke velocity in any age groups (*P* = 0.68) (Fig. 3D; Table A2). Initial maximal AP downstroke velocity increased from P5 to P21 by 2.06-fold in females (*P* < 0.001) and by 2.1-fold in males (*P* < 0.001). There were no sex differences in initial maximal AP downstroke velocity observed in any sex groups (*P* = 0.80) (Fig. 3E; Table A2). To assess changes in input processing capabilities, we next analyzed properties of the initial gain. Initial gain decreased from P5 to P21 by 1.9-fold in females (*P* < 0.001) and by 2.33-fold in males (*P* < 0.001). Sex-related differences appeared at P8 (*P* = 0.007) as female neurons showed higher initial gain (Fig. 3F; Table A2).

**Fig. 3.**
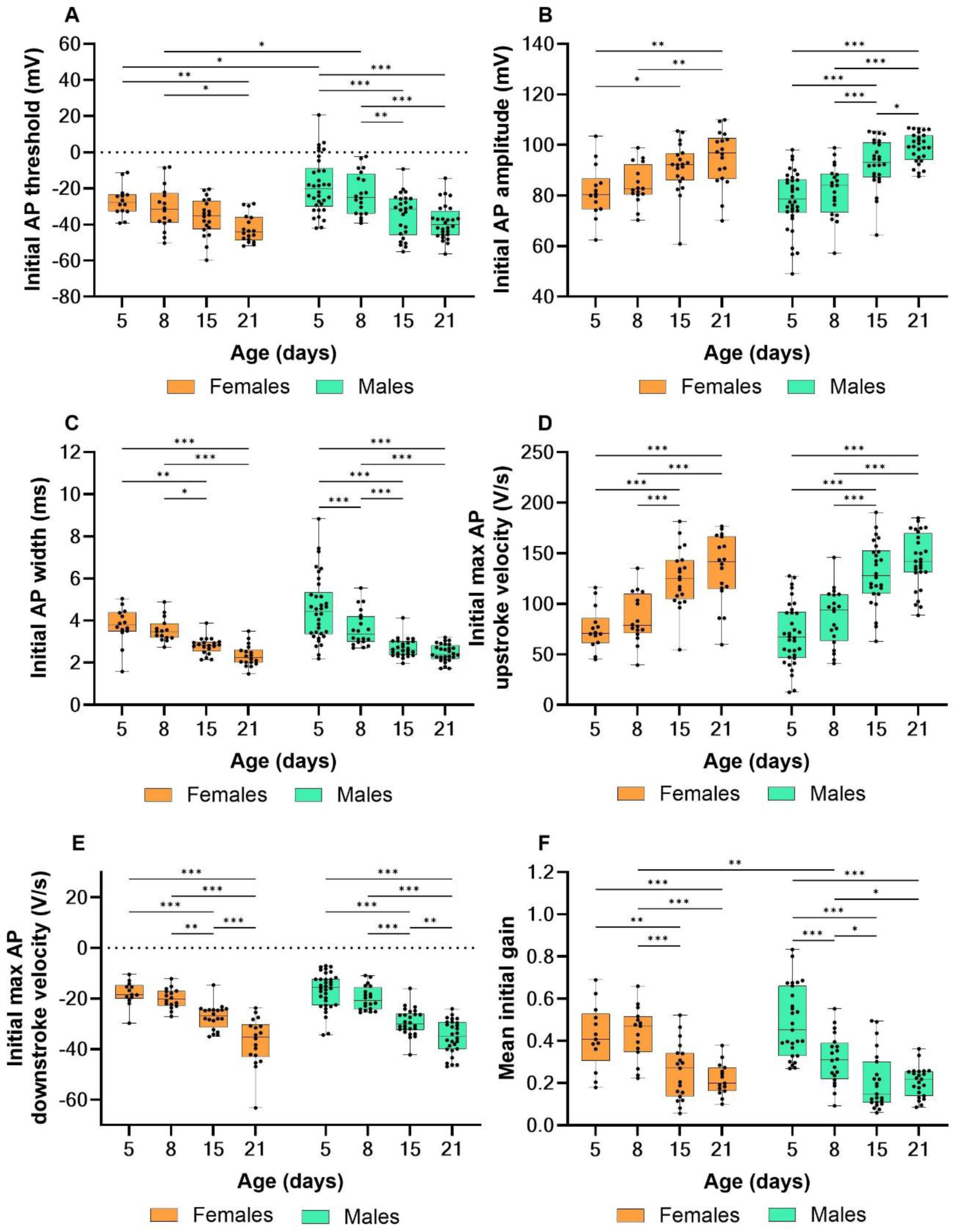
Changes in active initial electrophysiological properties during development. Developmental stages are shown as postnatal days (P5, P8, P15, P21). Asterisks indicate significant differences between groups (\**P*<0.05, \*\**P*<0.01, \*\*\**P*<0.001). Orange indicates females; green indicates males. (**A**) AP threshold became more hyperpolarized during the development, with sex differences at P5–P8. (**B**) AP amplitude increased during development, while (**C**) AP width decreased during development. (**D**) AP upstroke velocity increased with age. (**E**) AP downstroke velocity became faster by P21. (**F**) The slope of the initial f–I curve decreased with age, with a sex difference at P8.

### Steady AP electrophysiological properties

Steady AP threshold changed from P5 to P21, hyperpolarising by 1.61-fold in females (*P* = 0.001) and by 2.05-fold in males (*P* < 0.001). Sex differences were observed at P5 (*P* = 0.04) and P8 (*P* = 0.03), with a more depolarised steady AP threshold in males (Fig. 4A; Table A3). Steady AP amplitude increased from P5 to P21 by 23.2% in females (*P* < 0.001) and by 22.6% in males (*P* < 0.001). There were no significant sex differences observed in any age groups (*P* = 0.24) (Fig. 4B; Table A3). Steady AP width decreased from P5 to P21 by 2.11-fold in females (*P* < 0.001) and by 2.22-fold in males (*P* < 0.001). There were no sex differences in steady AP width found in any age groups (*P* = 0.20) (Fig. 4C; Table A3). Steady maximal AP upstroke velocity increased from P5 to P21 by 2.21-fold in females (*P* < 0.001) and by 2.11-fold in males (*P* < 0.001). There were no sex differences in steady maximal AP upstroke velocity found in any age groups (*P* = 0.69) (Fig. 4D; Table A3). Steady maximal AP downstroke velocity increased from P5 to P21 by 2.56-fold in females (*P* < 0.001) and by 2.39-fold in males (*P* < 0.001). Sex differences in steady maximal AP downstroke velocity were observed at P15 (higher in males, *P* = 0.03) and at P21 (higher in females, *P* = 0.02) (Fig. 4E; Table A3). Steady gain decreased from P5 to P21 by 2.83-fold in females (*P* < 0.001) and by 3-fold in males (*P* < 0.001). Significant differences between sex groups were observed at P5 (higher in males, *P* = 0.008) and P8 (higher in females, *P* < 0.001) (Fig. 4F; Table A3). Altogether, our results showed that during the first three postnatal weeks, both initial and steady APs increased in amplitude and decreased in duration, the threshold hyperpolarised, maximal upstroke and downstroke velocities increased, and initial and steady gain decreased in mouse hippocampal CA1 pyramidal neurons. These developmental changes in active electrophysiological properties could be linked to postnatal variations in the density and the kinetics of ionic channels that regulate the generation of action potentials.

**Fig. 4.**
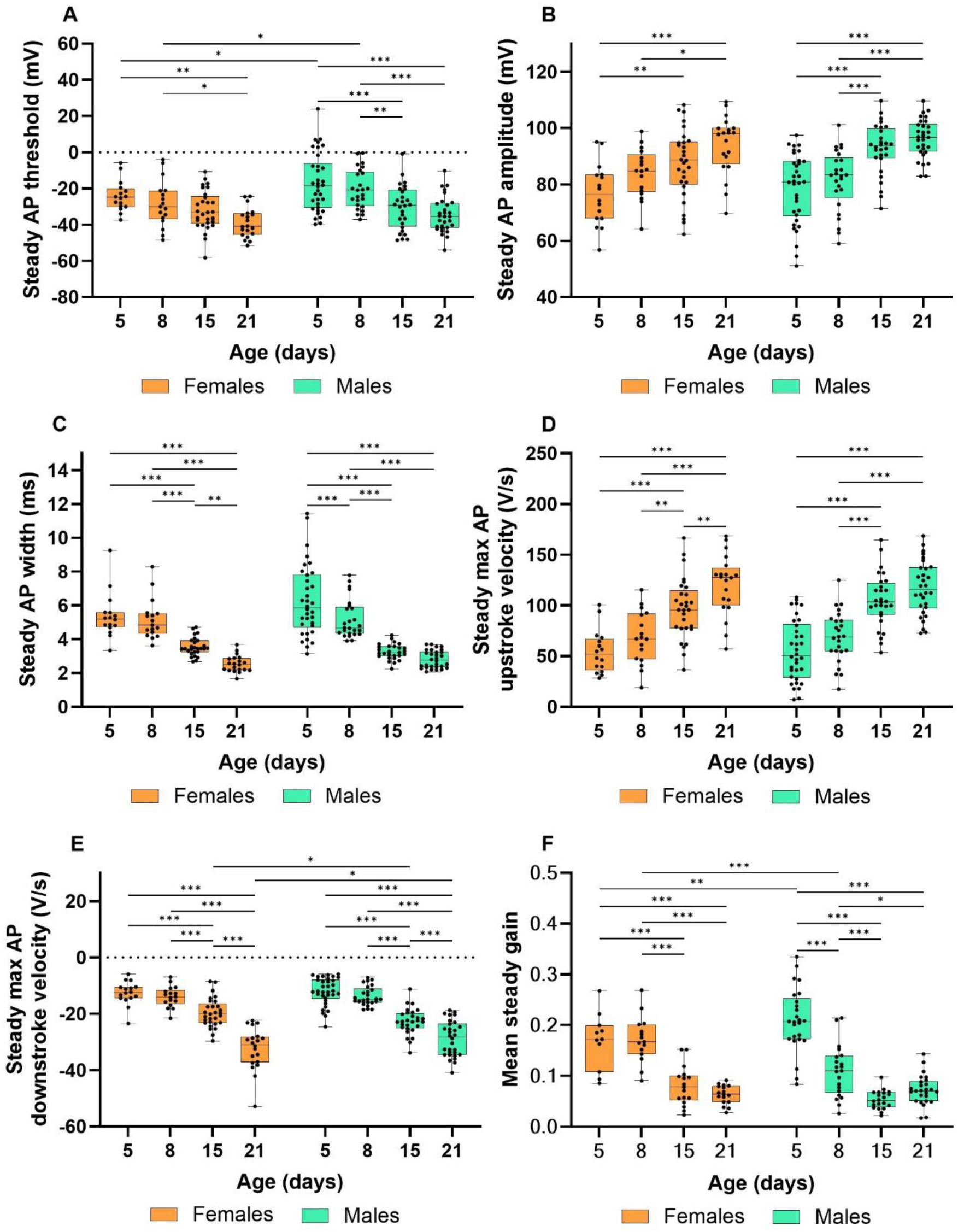
Changes in active steady electrophysiological properties during development. Developmental stages are shown as postnatal days (P5, P8, P15, P21). Asterisks indicate significant differences between groups (\**P*<0.05, \*\**P*<0.01, \*\*\**P*<0.001). Orange indicates females; green indicates males. (**A**) AP threshold became more hyperpolarized during the development, with sex differences at P5–P8. (**B**) AP amplitude increased during development, while (**C**) AP width decreased during development. (**D**) AP upstroke velocity increased with age. (**E**) AP downstroke velocity became faster by P21. (**F**) Initial gain decreased with age, with a sex difference at P5 and P8.

### The effect of spike-frequency adaptation on active electrophysiological properties

Spike-frequency adaptation reflects how the response of neurons as they process slowly changing inputs and can affect intrinsic electrophysiological properties; therefore, we examined how active electrophysiological properties are affected by adaptation. In all age and sex groups, the AP threshold depolarised during spike-frequency adaptation (*P* < 0.001) (Fig. 5A-C). In contrast, AP amplitude decreased only in the P21 male group (*P* < 0.001) (Fig. 5D-F). AP width increased during spike-frequency adaptation in all groups (*P* < 0.001), with a systematically larger increase at younger ages, suggesting that mature neurons are more resistant to adaptation-induced changes in AP width. (Fig. 5G-I). Spike-frequency adaptation affected maximal AP upstroke and downstroke velocities as well. They both decreased in all age and sex groups (*P* < 0.001) (Fig. 6A-F). Moreover, in both males and females in all age groups, the gain was significantly decreased during spike frequency adaptation (*P* < 0.001) (Fig. 7A-B).

**Fig. 5.**
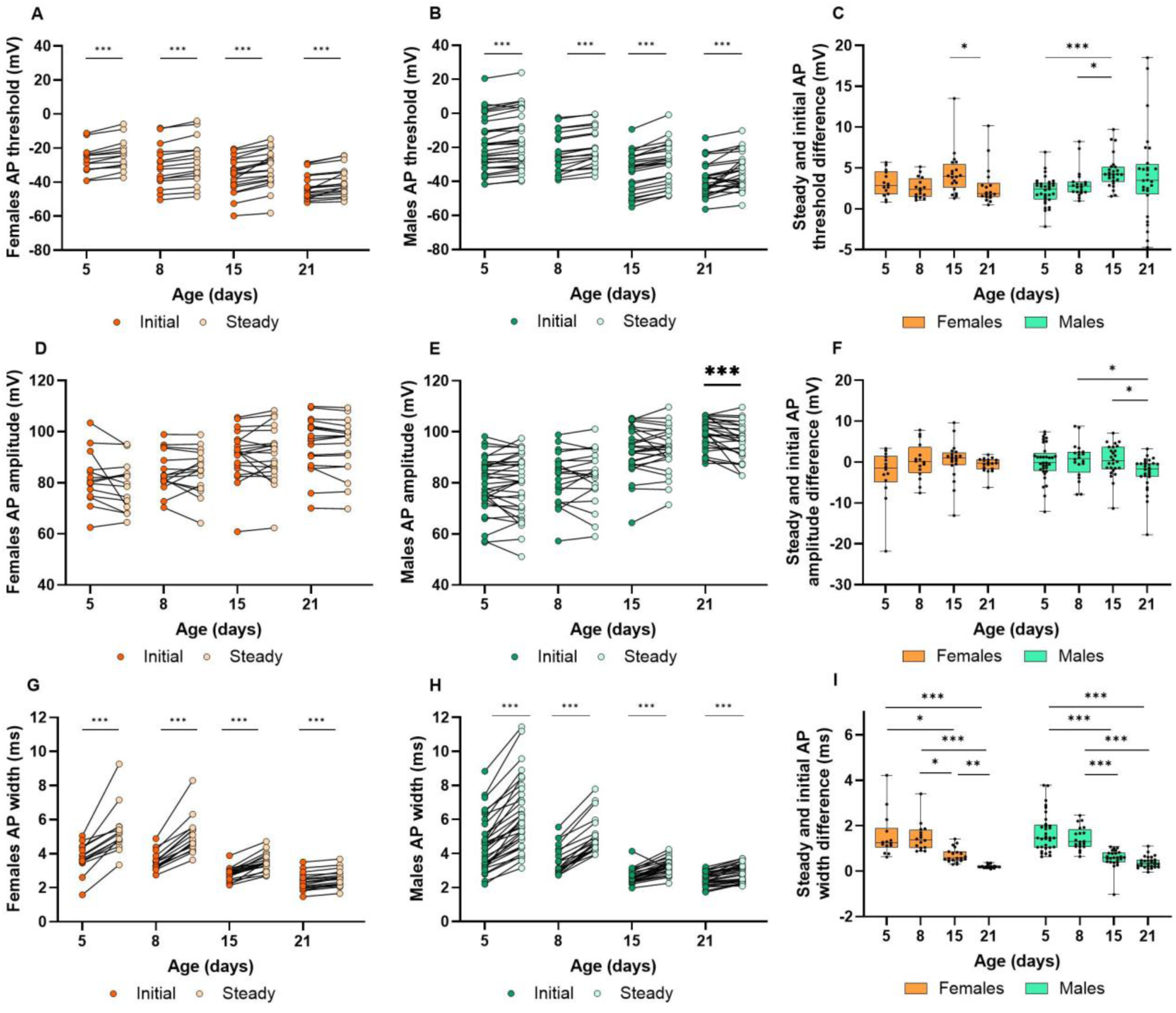
The effect of spike-frequency adaptation on AP threshold, AP amplitude and AP width. Asterisks indicate significant differences between groups (\**P*<0.05, \*\**P*<0.01, \*\*\**P*<0.001). Orange indicates females; green indicates males. Dark-coloured dots indicate initial AP properties, whereas light-coloured symbols indicate steady-state values after adaptation. (**A-C**) AP threshold depolarised during spike-frequency adaptation in both males and females in all age groups. (**D-F**) AP amplitude decreased only in the P21 male group. (**G-I**) AP width increased during spike-frequency adaptation in both males and females in all age groups.

**Fig. 6.**
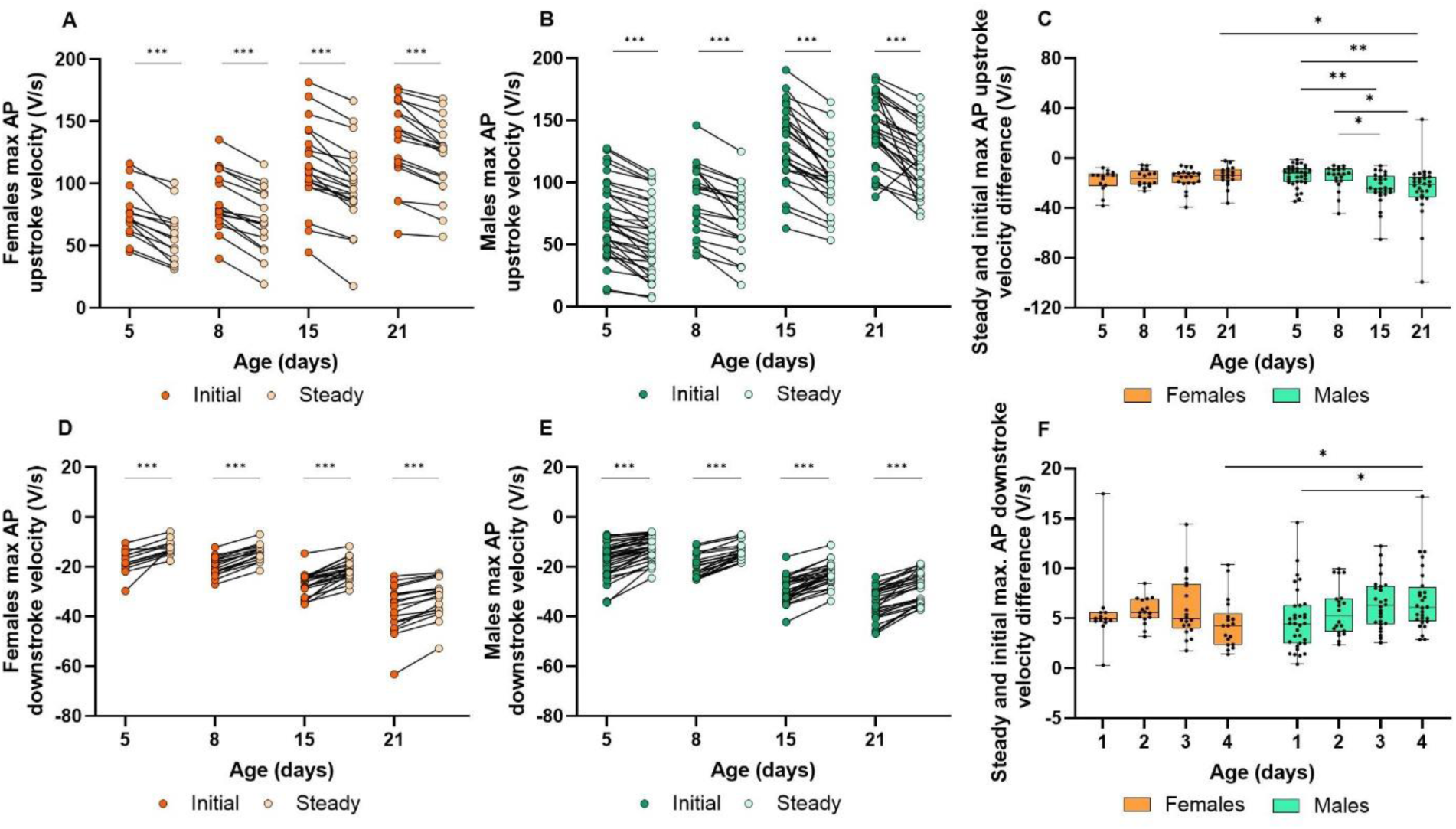
The effect of spike-frequency adaptation on AP maximal upstroke and downstroke velocities. Asterisks indicate significant differences between groups (\**P*<0.05, \*\**P*<0.01, \*\*\**P*<0.001). Orange indicates females; green indicates males. Dark-coloured dots indicate initial AP properties, whereas light-coloured symbols indicate steady-state values after adaptation. (**A-C**) Maximal AP upstroke and (**D-F**) maximal AP downstroke velocities decreased in all age and sex groups.

**Fig. 7.**
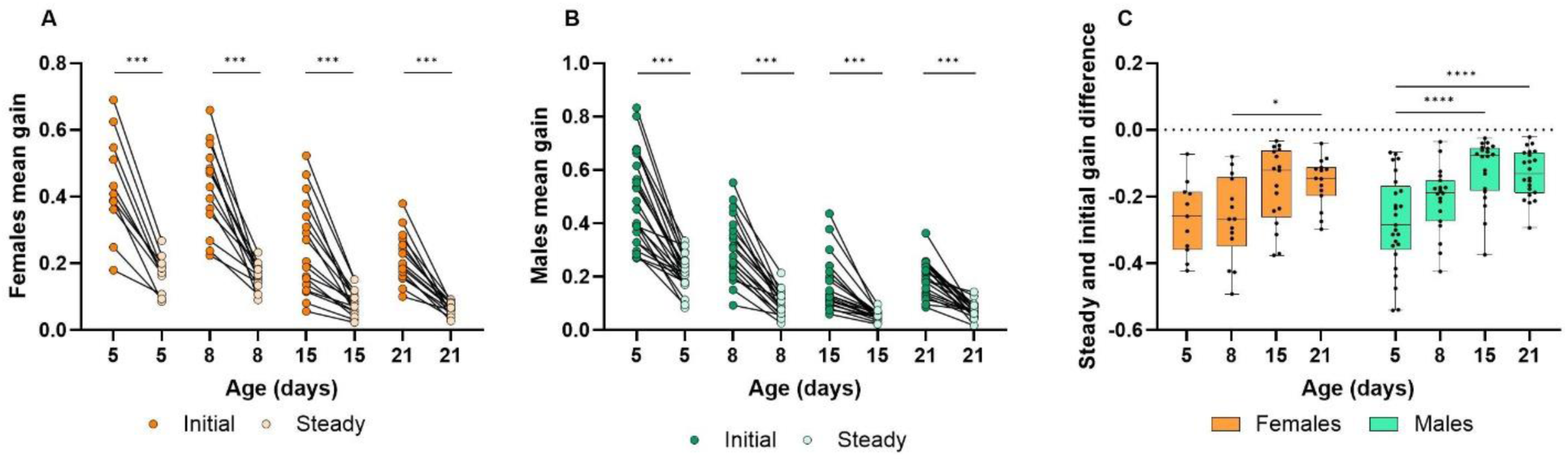
The effect of spike-frequency adaptation on gain. Asterisks indicate significant differences between groups (\**P*<0.05, \*\**P*<0.01, \*\*\**P*<0.001, \*\*\*\**P*<0.0001). Dark-coloured dots indicate initial gain, whereas light-coloured symbols indicate steady-state gain after adaptation. (**A-C**) The gain was significantly decreased during spike frequency adaptation in both males and females at all ages.

Spike-frequency adaptation resulted in changes in active electrophysiological properties, including increased action potential width, depolarization of threshold, reduced maximal upstroke and downstroke velocities, and decreased gain, leading to decreased excitability. Most of these adaptation-related changes were similar across ages and sexes; however, action potential width during spike-frequency adaptation showed a clear age dependence, with a progressively smaller increase observed in more mature neurons.

### The development of synaptic activity

The maturation of intrinsic properties of CA1 neurons coincides with the period of proliferation and refinement of synaptic connections (Yang et al., 2024; Harris et. al., 1992; Yasuda et al., 2011; Pokorný and Yamamoto, 1981). Changes in neuronal firing may affect network activity, which we evaluated by examining synaptic parameters, such as the peak amplitude and the duration of the interevent interval of spontaneous excitatory postsynaptic currents (sEPSC). sEPSC interevent intervals, characterizing synaptic connectivity between CA3–CA1 pyramidal neurons, decreased by 5.57-fold from P5 to P21 in females (*P* < 0.001) and by 2.37-fold in males (*P* < 0.001). Sex related differences were observed at P5 (*P* < 0.001) and P8 (*P* < 0.001) as female neurons showed longer sEPSC interevent intervals (Fig. 8A; Table A4). The developmental decrease in sEPSC interevent intervals reflects an increase in synaptic event frequency, suggesting increased functional synaptic connectivity between CA3– CA1 pyramidal neurons. To characterize the strength of synaptic connectivity, peak amplitudes of sEPSC were evaluated throughout postnatal development. sEPSC peak amplitude did not change in both males and females during the development. There were no significant sex related differences in sEPSC peak amplitude (*P* = 0.11) (Fig. 8B; Table A4). Although mean sEPSC amplitude did not differ significantly between developmental stages, distribution-based analysis was performed to detect potential age-dependent shifts in sEPSC amplitude that may not be reflected by group averages (Fig. A1). In both females and males, normalized histograms and cumulative probability plots revealed almost overlapping amplitude distributions between P5 and P21, indicating minimal age-dependent changes in synaptic strength despite Kolmogorov-Smirnov (KS) test significance (*P* < 0.001) driven by large event numbers (Fig. A1 A–D). Although no sex differences were detected when comparing mean sEPSC peak amplitudes, distribution-based analyses revealed sex-dependent shifts in sEPSC amplitude distributions at early postnatal stages. At P5 and P8, males show a higher number of smaller-amplitude events compared to females (KS test *P* < 0.001; Fig. A2 A–D). In contrast, at P15 and P21 the amplitude distributions largely overlapped, indicating minimal biological differences between sexes despite statistical significance detected by the KS test (Fig. A3 A–D). Together, these results indicate that developmental changes in synaptic connectivity between CA3–CA1 pyramidal neurons are primarily driven by an increase in synaptic event frequency, whereas synaptic strength, assessed by sEPSC amplitudes, remains largely stable across age and sex.

**Fig. 8.**
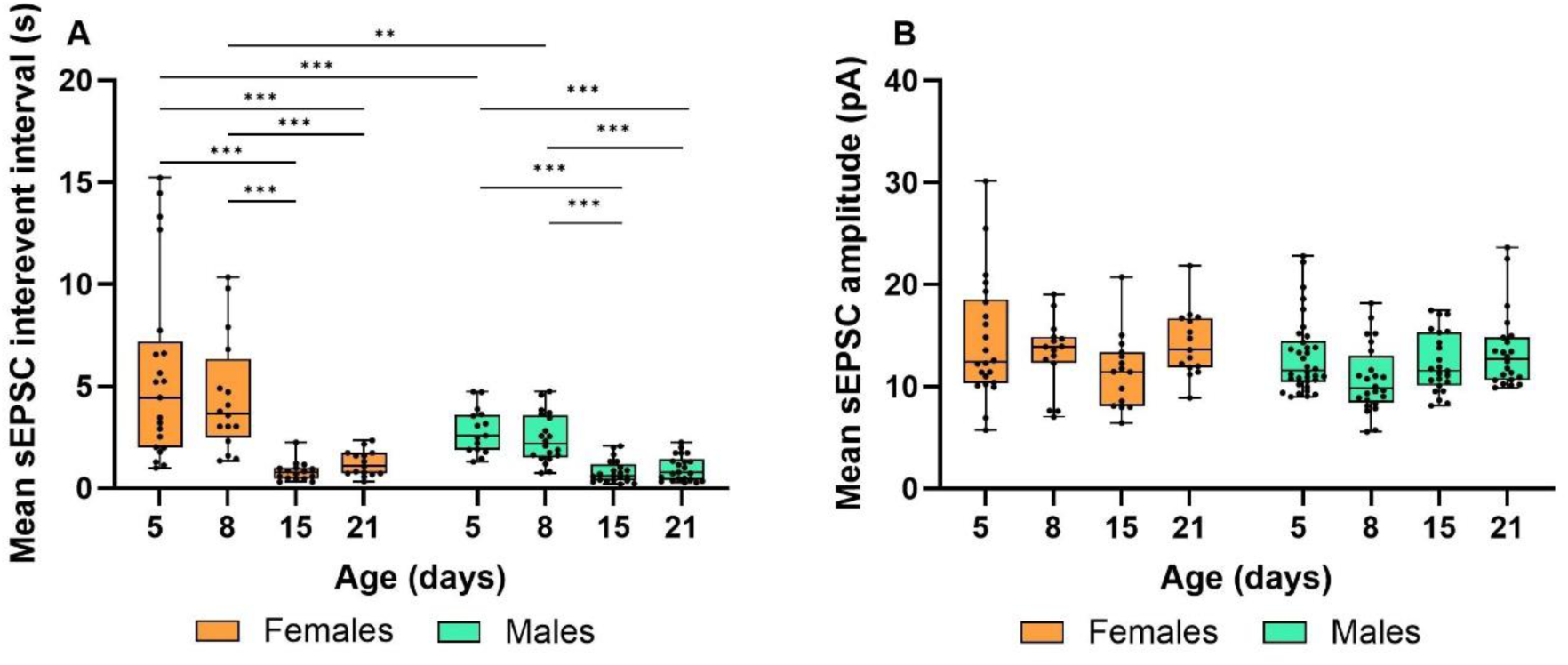
Changes in synaptic activity during development in females and males. Developmental stages are shown as postnatal days (P5, P8, P15, P21). Asterisks indicate significant differences between groups (\**P*<0.05, \*\**P*<0.01, \*\*\**P*<0.001). Orange indicates females and green indicates males. (**A**) Mean sEPSC interevent intervals decreased with age both in males and females, with sex differences observed at P5 and P8. (**B**) There are no differences in mean peak sEPSC amplitude during the development in both sexes. There were no sex differences observed as well.

## Discussion

In this study, we investigated the maturation of hippocampal CA1 pyramidal neurons in mice, characterizing intrinsic membrane and firing properties, as well as synaptic activity throughout postnatal development. We found that the intrinsic neuronal characteristics and firing properties of mouse hippocampal CA1 pyramidal neurons transformed substantially from the 5th to the 21st day of postnatal development. The input resistance and time membrane constant decreased, while membrane capacitance increased.

Action potential amplitude increased, width decreased, threshold hyperpolarized, and maximum upstroke and downstroke velocities increased. The gain decreased in both male and female mice from P5 to P21 as well. Moreover, the interevent interval of recorded sEPSC decreased from P5 to P21, while the amplitude remained unchanged.

Despite the electrophysiological changes observed during the first three weeks of postnatal development, the most significant alterations in active membrane properties and synaptic activity were determined during the second postnatal week (P8 to P15), indicating a critical period for rapid formation, plasticity, and maturation of the mouse hippocampus circuitry. This could be associated with the period of eye opening between P12 and P15, after which rodent pups first begin to explore their environment (Ciganok-Hückels et al., 2023; Hoy and Niell, 2015; Wills et al., 2010; Lu and Constantine-Paton, 2004). Moreover, the major changes during the second postnatal week might be related to the peak in synaptogenesis of CA1 neurons occurring from two to four weeks postnatally (Yang et al., 2024; Steward and Falk, 1991) and increasing microglia activity during the second postnatal week as well (Weinhard et al., 2018; Pagani et al., 2015; Paolicelli et al., 2011). However, the passive membrane properties changed substantially during the first postnatal week (from P5 to P8), suggesting that alterations in neuronal membrane could be associated with the maturation of active neuronal properties. Other studies have shown that, during postnatal development, the input resistance and time constant of other types of neurons in rats and mice decrease (Brahimi et al., 2023; Kroon et al., 2019; McCormick and Prince, 1987), in line with our observations. The changes of active membrane properties identified in this study were similar to the findings of other authors, who indicated that in rat cerebral cortex and hippocampal CA1 pyramidal neurons, AP threshold hyperpolarizes, width (duration) decreases, amplitude, upstroke and downstroke velocities increase (Brahimi et al., 2023; Sánchez-Aguilera et al., 2020; Giglio and Storm, 2014; Sengupta et al., 2013; Spigelman et al., 1992). The same tendencies in AP threshold, width, and amplitude changes were observed in mouse medial prefrontal cortex pyramidal neurons as well (Kroon et al., 2019). Based on our results, we hypothesize that hippocampal CA1 pyramidal neurons undergo changes in AP parameters due to the increasing number of ion channels in the membrane. Moreover, our results confirm that these developmental changes are similar in rats and mice, despite their anatomical differences. To investigate the firing characteristics of neurons, we analyzed the gain of the f-I relation, a parameter of neuronal excitability, which describes how a neuron’s firing rate varies with injected current. Our results revealed that both the initial and steady gains decreased during postnatal development of hippocampal CA1 neurons. Meanwhile, other electrophysiological studies of the mouse cortical regions have demonstrated that neural gain remains consistent across different ages (Ciganok-Hückels et al., 2023; Kinnischtzke et al., 2012). The decrease in neural gain in CA1 neurons may indicate unique maturation processes in the hippocampus that result in more precise regulation of neuronal firing. Our findings also indicated that spike-frequency adaptation caused fatigue of neuronal output and affected the firing properties of pyramidal neurons, resulting in an increase in action potential width, a decrease in maximal upstroke and downstroke velocities, a depolarization of the threshold, and a decrease in gain. These results might indicate the important role of spike-frequency adaptation in suppressing a neuron’s response to a constant stimulus enabling neuron to quickly detect changes in stimuli and efficiently process information (Adibi et al., 2013). Anatomical and immunohistochemical studies have shown that the number of synapses between CA3 and CA1 pyramidal neurons increases during postnatal development, suggesting enhanced circuit connectivity (Yang et al., 2024; Paolicelli et al., 2011; Hsia et al., 1998)(Harris et al., 1992). Consistent with this structural maturation, our electrophysiological analysis revealed a decrease in sEPSC interevent intervals, while sEPSC amplitudes remained stable across development. At P21, the sEPSC interevent intervals in our recordings (∼1.2 – 1.4 s) were comparable to values reported for similarly aged hippocampal pyramidal neurons (∼0.8 – 1.2 s at P20–26; Baxter et al., 2011; Lu et al., 2018). At earlier stages, intervals at P5 (∼3.1 – 5.5 s) were of similar values to those reported in neonatal mice (∼4.3 s at P0–2; Orav et al., 2019). Notably, these values are typically reported as control data in studies not primarily focused on developmental trajectories. Thus, our data at P5 and intermediate ages up to P21 provide additional points along the postnatal developmental trajectory. Together, these findings could suggest that the developmental increase in excitatory synaptic activity primarily reflects an increase in the number of functional CA3–CA1 synapses, rather than changes in the strength of individual synaptic events. In parallel with functional synaptic maturation, postnatal hippocampal development is accompanied by dynamic microglial activity, which is known to contribute to synapse formation and elimination (Miyamoto et al., 2016; Sipe et al., 2016; Zhan et al., 2014; Parkhurst et al., 2013;

Schafer et al., 2012; Paolicelli et al., 2011) (Andoh and Ryuta 2021). Microglial volume and phagocytic capacity increase during the second postnatal week, overlapping with periods of synaptic reorganization in the hippocampus (Weinhard et al., 2018).

Moreover, sex-dependent differences in hippocampal development have been widely reported. Due to the influence of sex hormones, dendritic spine density of hippocampal CA1 pyramidal neurons changes during maturation in males and varies across the estrous cycle in females, with higher spine density during proestrus compared to estrus (Gould et al., 1990; Woolley et al., 1990; Meyer et al., 1978). In addition, microglial volume and phagocytic capacity peak earlier in female mice than in males (Weinhard et al., 2018), and the spatial and temporal expression of androgen and estrogen receptors in the mouse brain differs between sexes (Mogi et al., 2015). The morphology of hippocampal CA3 pyramidal neurons is also known to differ between sexes (Hashimoto et al., 2023; Juraska et al., 1989). Importantly, many disorders associated with hippocampal dysfunction, such as epilepsy, schizophrenia, autism spectrum disorder, depression, and anxiety, differ between the sexes in terms of prevalence and/or presentation (Kight and McCarthy, 2020). Women are known to have a greater prevalence and cognitive decline in such diseases as depression and Alzheimer’s disease, and men show more severe cognitive impairments when having schizophrenia or Parkinson’s disease (Yagi and Galea, 2019). Based on these observations, it is crucial to consider sex as a factor in studies of hippocampal circuit maturation.

Therefore, although sex was not the primary focus of the present study, we examined male and female animals separately to assess potential sex-dependent differences in electrophysiological properties and synaptic activity during postnatal development.

During the first postnatal week, male neurons exhibited higher input resistance at P5, a more depolarized resting membrane potential at P8, and more depolarized action potential threshold values at both P5 and P8 compared to females. Sex-dependent differences in synaptic activity were also evident at early developmental stages, with females showing longer sEPSC inter-event intervals than males at P5 and P8; these differences were no longer detectable at later stages. Distribution-based analyses revealed sex-dependent shifts in sEPSC amplitude distributions at early postnatal stages. At P5 and P8, males show a higher number of smaller-amplitude events compared to females, while mature neurons showed overlapped amplitude distributions, indicating minimal biological differences between sexes.

## Conclusions

Overall, our study confirms significant postnatal developmental changes in the electrophysiological profiles of mouse hippocampal pyramidal neurons and provides a baseline for future electrophysiological studies of postnatal hippocampal maturation in mice. The transformation of neuronal firing activity results in faster responses to a stimulus, the generation of action potentials of higher amplitude and shorter duration, and more precise control of neuronal firing with age. Simultaneously, alterations in synaptic activity emerge, revealing increased functional connectivity during postnatal development. Moreover, we observed the influence of sex on the trajectory of development only, as the sex differences in electrophysiological properties that occurred during the first postnatal week did not persist further into the second and third postnatal weeks. We determined that the first postnatal week is a critical period for the development of passive membrane properties, which could be largely attributed to the increase in neuronal size, arborization, and incorporation of diverse membrane channels. However, we found the second postnatal week to be critical for the maturation of active membrane properties and for changes in functional hippocampus connectivity. Despite changes during critical periods, neurons are known to maintain homeostatic plasticity, adjusting their intrinsic and firing properties in response to increasing excitatory synaptic activity (Turrigiano and Nelson, 2004). However, the homeostatic mechanisms and the relationship between neuronal network activity and membrane properties remain unknown. Altogether, the developmental changes in electrophysiological profiles of CA1 pyramidal neurons may contribute to shaping hippocampal circuitry and lead to more effective information processing in the mature brain.

## Appendix

**Table A1.**
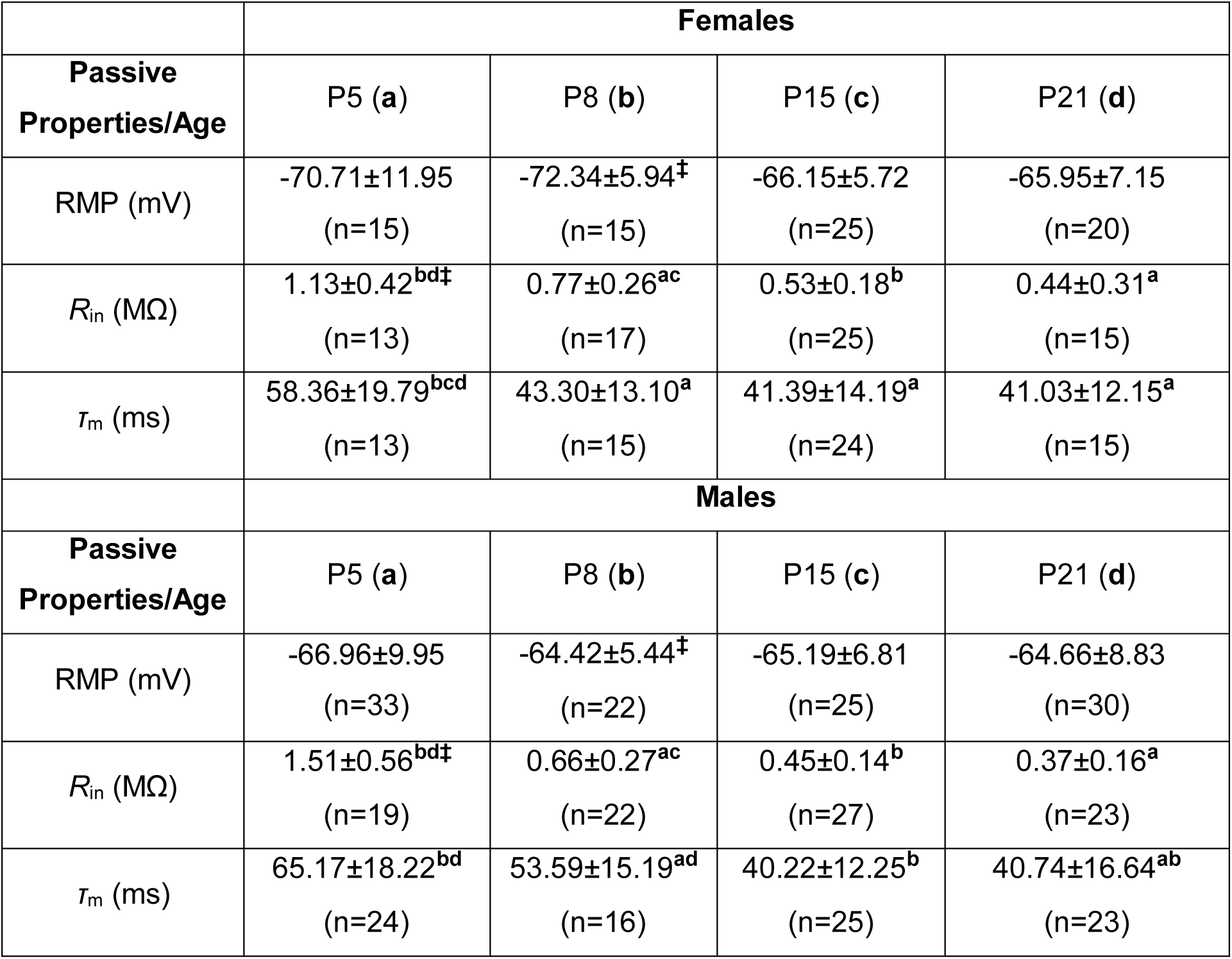
Resting membrane potential (RMP), input resistance (Rin), membrane time constant (τc) and membrane capacitance (Cm) during the development in females and males. Age groups are assigned fixed labels (P5 = **a**, P8 = **b**, P15 = **c**, P21 = **d**). Superscript letters next to each value indicate the specific age groups from which they differ significantly. For example, membrane capacitance at P21 in females is labelled “abc”, indicating that it differs significantly from P5 (**a**), P8 (**b**), and P15 (**c**). Sex differences within the same age are marked with **‡**.

|  | **Females** |  |  |  |
| --- | --- | --- | --- | --- |
| **Passive Properties/Age** | **P5 (a)** | **P8 (b)** | **P15 (c)** | **P21 (d)** |
| RMP (mV) | -70.71±11.95<br>(n=15) | -72.34±5.94<sup>‡</sup><br>(n=15) | -66.15±5.72<br>(n=25) | -65.95±7.15<br>(n=20) |
| $R_{in}$ (MΩ) | 1.13±0.42<sup>bd‡</sup><br>(n=13) | 0.77±0.26<sup>ac</sup><br>(n=17) | 0.53±0.18<sup>b</sup><br>(n=25) | 0.44±0.31<sup>a</sup><br>(n=15) |
| $\tau_m$ (ms) | 58.36±19.79<sup>bcd</sup><br>(n=13) | 43.30±13.10<sup>a</sup><br>(n=15) | 41.39±14.19<sup>a</sup><br>(n=24) | 41.03±12.15<sup>a</sup><br>(n=15) |
|  | **Males** |  |  |  |
| **Passive Properties/Age** | **P5 (a)** | **P8 (b)** | **P15 (c)** | **P21 (d)** |
| RMP (mV) | -66.96±9.95<br>(n=33) | -64.42±5.44<sup>‡</sup><br>(n=22) | -65.19±6.81<br>(n=25) | -64.66±8.83<br>(n=30) |
| $R_{in}$ (MΩ) | 1.51±0.56<sup>bd‡</sup><br>(n=19) | 0.66±0.27<sup>ac</sup><br>(n=22) | 0.45±0.14<sup>b</sup><br>(n=27) | 0.37±0.16<sup>a</sup><br>(n=23) |
| $\tau_m$ (ms) | 65.17±18.22<sup>bd</sup><br>(n=24) | 53.59±15.19<sup>ad</sup><br>(n=16) | 40.22±12.25<sup>b</sup><br>(n=25) | 40.74±16.64<sup>ab</sup><br>(n=23) |

**Table A2.**
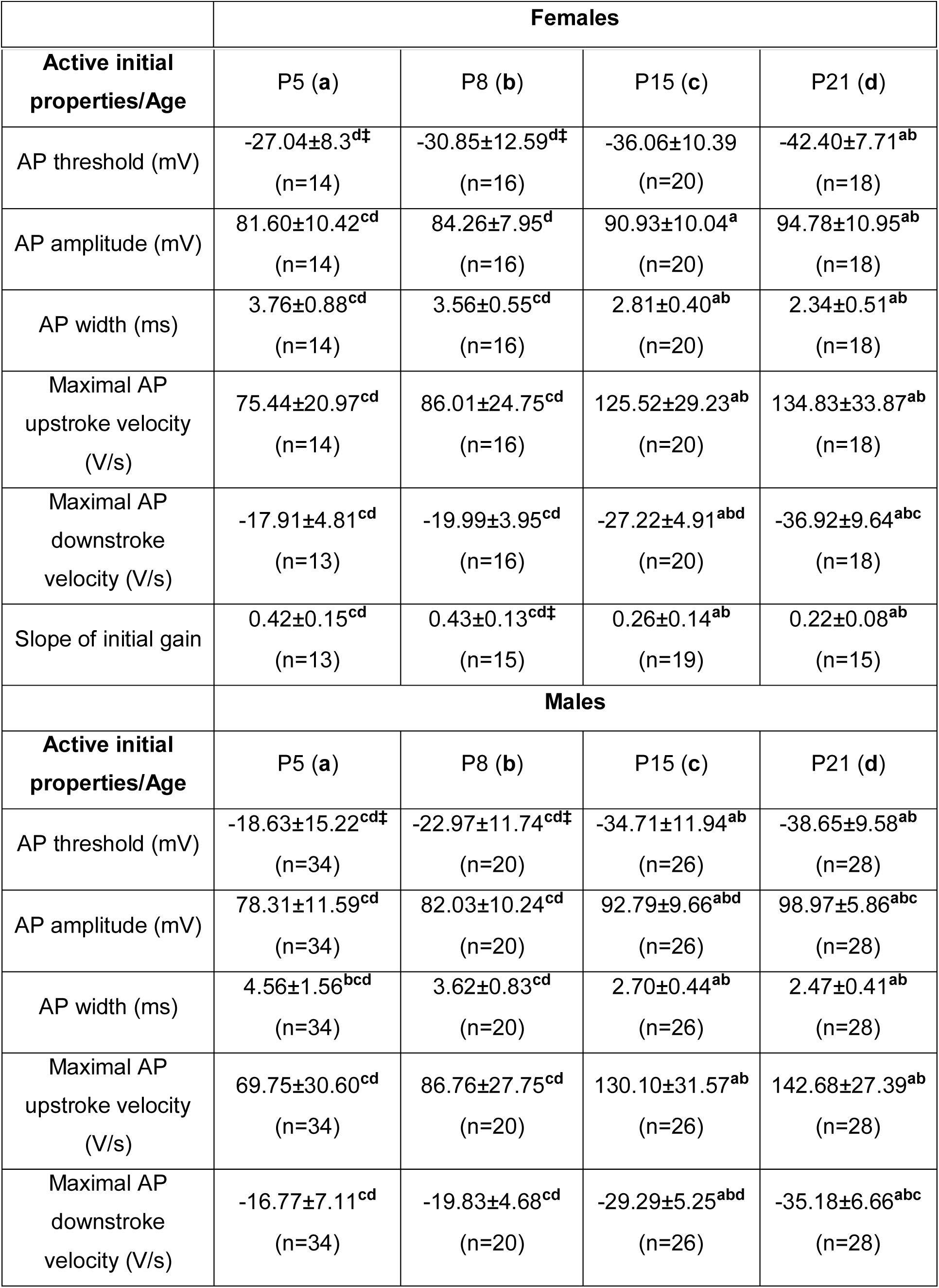
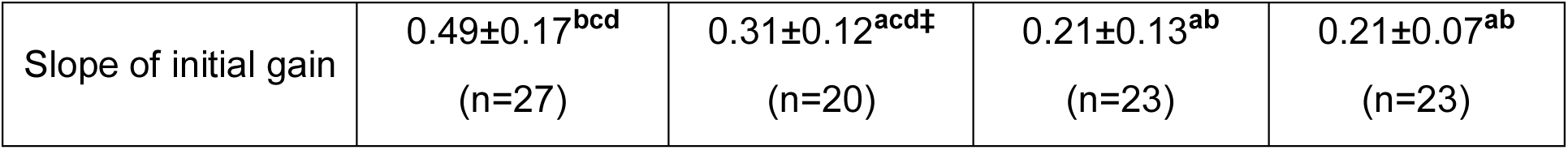
AP threshold (mV), AP amplitude (mV), AP width (ms), maximal AP upstroke/downstroke velocities (V/s) and slope of initial gain during the development in females and males. Age groups are assigned fixed labels (P5 = **a**, P8 = **b**, P15 = **c**, P21 = **d**). Superscript letters next to each value indicate the specific age groups from which they differ significantly. For example, AP amplitude at P21 in males is labelled “**abc**”, indicating that it differs significantly from P5 (**a**), P8 (**b**), and P15 (**c**). Sex differences within the same age are marked with **‡**.

**Table A3.** Steady-state AP threshold (mV), AP amplitude (mV), AP width (ms), maximal AP upstroke/downstroke velocities (V/s) and slope of steady gain during the development in females and males. Age groups are assigned fixed labels (P5 = **a**, P8 = **b**, P15 = **c**, P21 = **d**). Superscript letters next to each value indicate the specific age groups from which they differ significantly. For example, AP width at P21 in females is labelled “**abc**”, indicating that it differs significantly from P5 (**a**), P8 (**b**), and P15 (**c**). Sex differences within the same age are marked with **‡**.

|  | **Females** |  |  |  |
| --- | --- | --- | --- | --- |
| **Active steady properties/Age** | P5 (**a**) | P8 (**b**) | P15 (**c**) | P21 (**d**) |
| AP threshold (mV) | -24.17±8.71<sup>d‡</sup><br>(n=15) | -28.36±12.82<sup>d‡</sup><br>(n=17) | -32.09±10.84<br>(n=29) | -39.08±8.08<sup>ab</sup><br>(n=20) |
| AP amplitude (mV) | 77.75±9.88<sup>cd</sup><br>(n=15) | 83.99±8.79<sup>d</sup><br>(n=17) | 87.41±12.25<sup>a</sup><br>(n=29) | 94.13±10.34<sup>ab</sup><br>(n=20) |
| AP width (ms) | 5.33±1.36<sup>cd</sup><br>(n=15) | 5.18±1.20<sup>cd</sup><br>(n=17) | 3.55±0.53<sup>abd</sup><br>(n=27) | 2.54±0.49<sup>abc</sup><br>(n=20) |
| Maximal AP upstroke velocity (V/s) | 56.26±21.01<sup>cd</sup><br>(n=15) | 68.33±26.00<sup>cd</sup><br>(n=17) | 96.87±29.62<sup>abd</sup><br>(n=29) | 120.69±29.56<sup>abc</sup><br>(n=20) |
| Maximal AP downstroke velocity (V/s) | -12.94±4.10<sup>cd</sup><br>(n=15) | -13.93±3.66<sup>cd</sup><br>(n=17) | 19.66±5.11<sup>abd‡</sup><br>(n=29) | -32.32±7.47<sup>abc‡</sup><br>(n=20) |
| Slope of steady gain | 0.17±0.06<sup>cd‡</sup><br>(n=11) | 0.17±0.05<sup>cd‡</sup><br>(n=15) | 0.08±0.04<sup>ab</sup><br>(n=17) | 0.06±0.02<sup>ab</sup><br>(n=17) |
|  | **Males** |  |  |  |
| **Active steady properties/Age** | P5 (**a**) | P8 (**b**) | P15 (**c**) | P21 (**d**) |
| AP threshold (mV) | -16.83±16.05<sup>cd‡</sup><br>(n=34) | -20.43±10.86<sup>cd‡</sup><br>(n=25) | -30.42±12.06<sup>ab</sup><br>(n=28) | -34.55±9.77<sup>ab</sup><br>(n=30) |
| AP amplitude (mV) | 78.57±12.35<sup>cd</sup><br>(n=34) | 81.59±10.73<sup>cd</sup><br>(n=25) | 92.60±9.09<sup>ab</sup><br>(n=28) | 96.34±6.84<sup>ab</sup><br>(n=30) |
| AP width (ms) | 6.28±2.10<sup>bcd</sup><br>(n=34) | 5.18±1.14<sup>acd</sup><br>(n=25) | 3.27±0.46<sup>ab</sup><br>(n=28) | 2.83±0.52<sup>ab</sup><br>(n=30) |
| Maximal AP upstroke velocity (V/s) | 55.45±30.16<sup>cd</sup><br>(n=34) | 68.84±25.24<sup>cd</sup><br>(n=25) | 105.24±26.32<sup>bd</sup><br>(n=28) | 117.21±26.39<sup>ab</sup><br>(n=30) |
| Maximal AP downstroke velocity (V/s) | -12.04±4.80<sup>cd</sup><br>(n=34) | -13.70±3.42<sup>cd</sup><br>(n=25) | 22.71±4.61<sup>abd‡</sup><br>(n=28) | -28.81±6.26<sup>abc‡</sup><br>(n=30) |
| Slope of steady gain | 0.21±0.06<sup>bcd‡</sup><br>(n=25) | 0.11±0.05<sup>acd‡</sup><br>(n=21) | 0.05±0.02<sup>ab</sup><br>(n=20) | 0.07±0.03<sup>ab</sup><br>(n=26) |

**Table A4.**
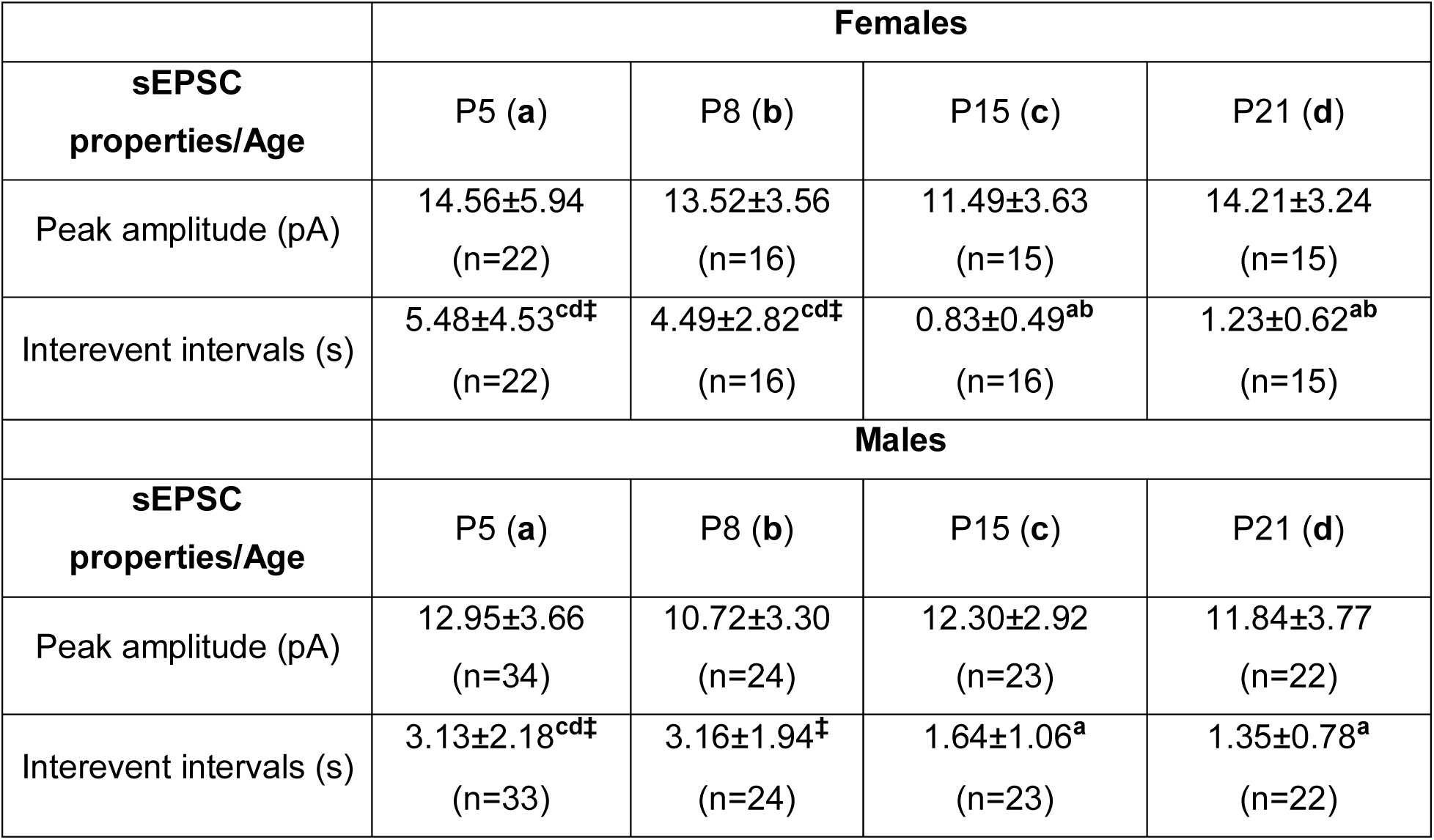
Peak sEPSC amplitude (pA) and sEPSC interevent intervals (s) during the development in females and males. Age groups are assigned fixed labels (P5 = **a**, P8 = **b**, P15 = **c**, P21 = **d**). Superscript letters next to each value indicate the specific age groups from which they differ significantly. For example, the value of sEPSC interevent intervals at P15 in females is labelled “**ab**”, indicating that it differs significantly from P5 (**a**), P8 (**b**). Sex differences within the same age are marked with **‡**.

**Fig. A1.**
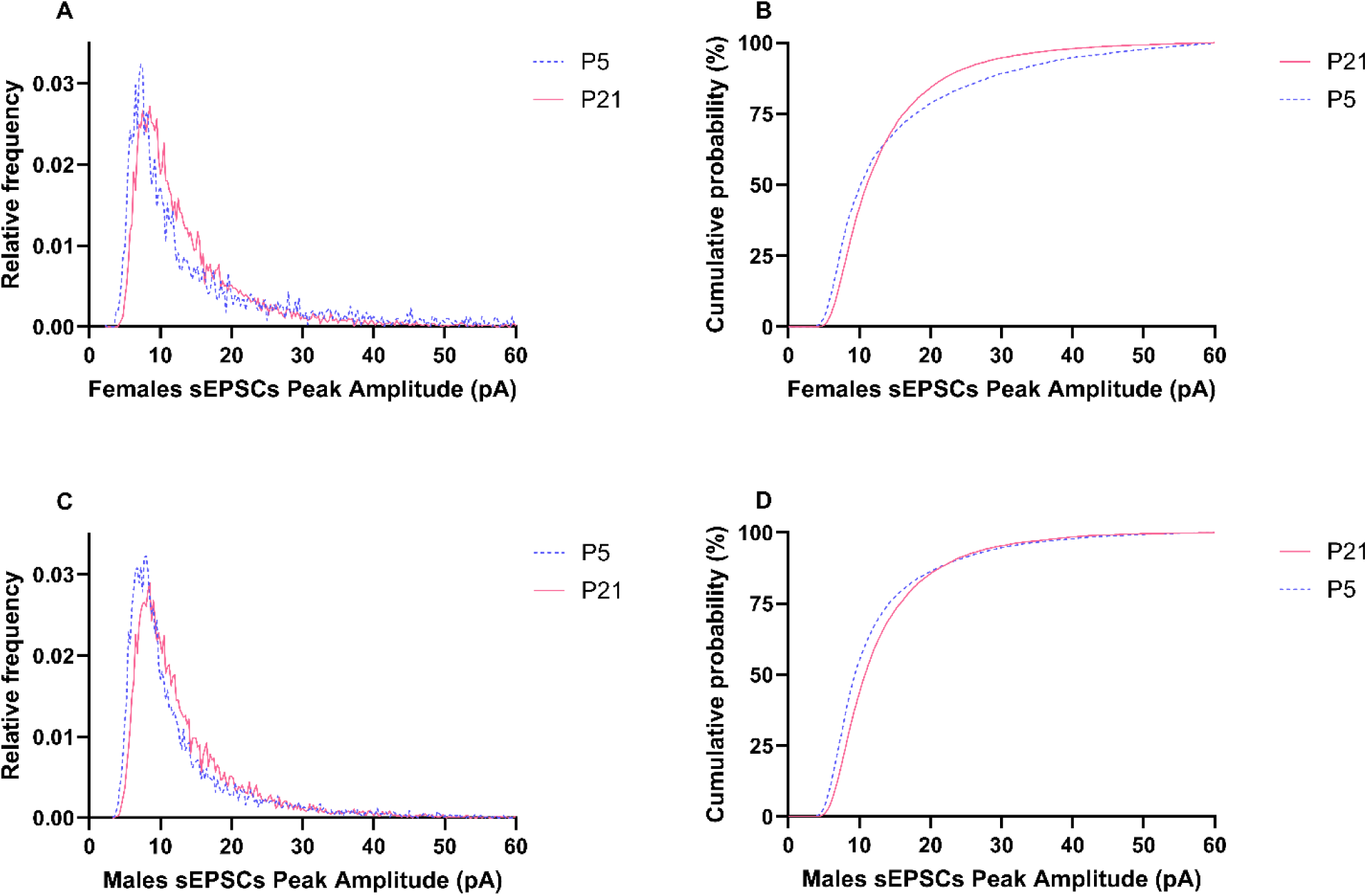
Distribution of sEPSC peak amplitudes during the development from P5 to P21. Normalized histograms illustrate the distributions of sEPSC peak amplitudes at P5 (blue/dashed) and P21 (red/solid) in females (**A**) and males (**C**). Corresponding cumulative probability plots compare sEPSC peak amplitude distributions in the early (P5) and late (P21) postnatal stages in females (**B**) and males (**D**).

**Fig. A2.**
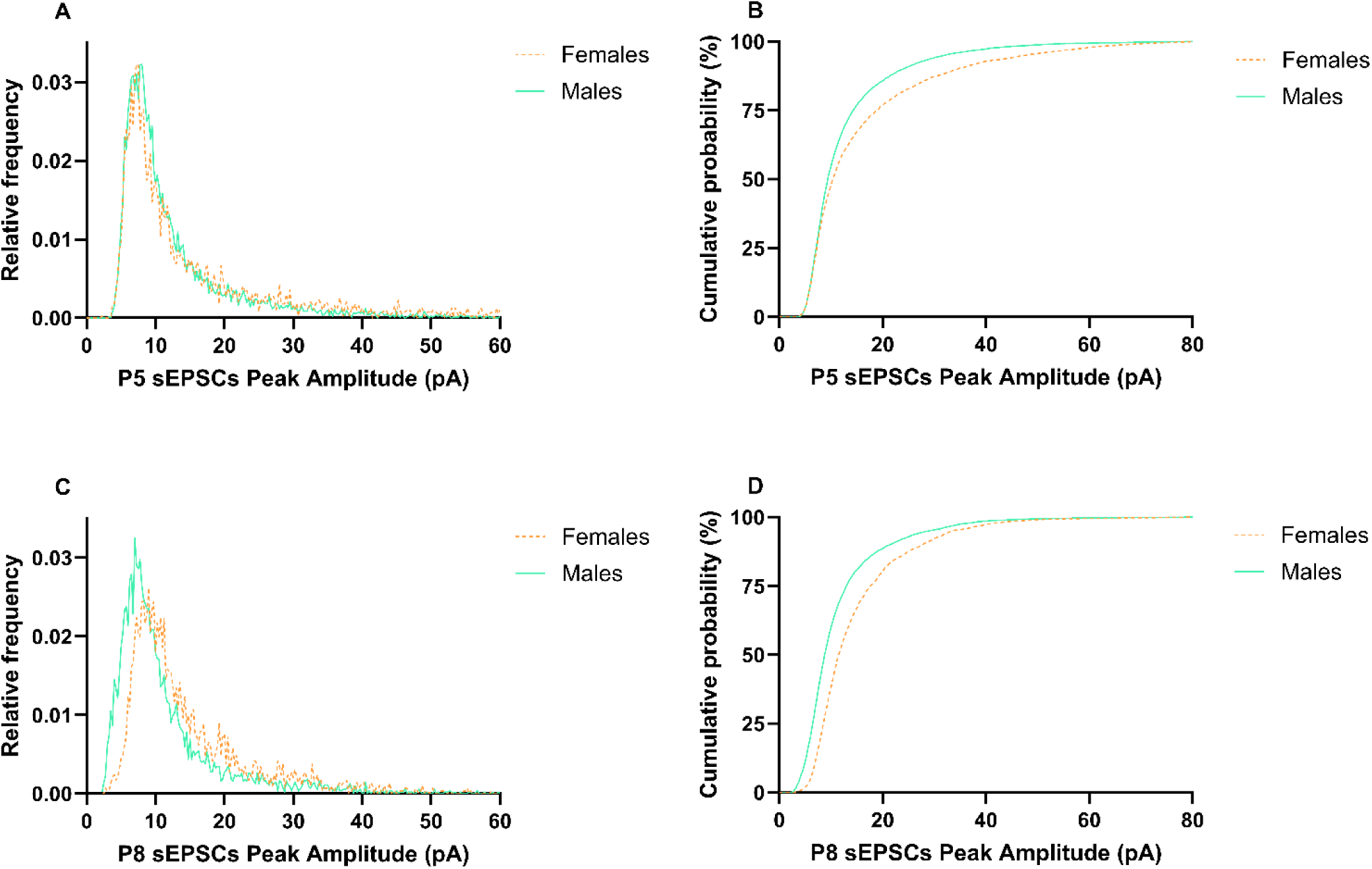
Distribution of sEPSC peak amplitudes at different early postnatal stages between sexes. Normalized histograms show the distributions of sEPSC peak amplitudes in females (orange/dashed) and males (green/solid) at P5 (**A**) and P8 (**C**). (**B, D**) Corresponding cumulative probability plots show the distributions of sEPSC peak amplitudes across sexes.

**Fig. A3.**
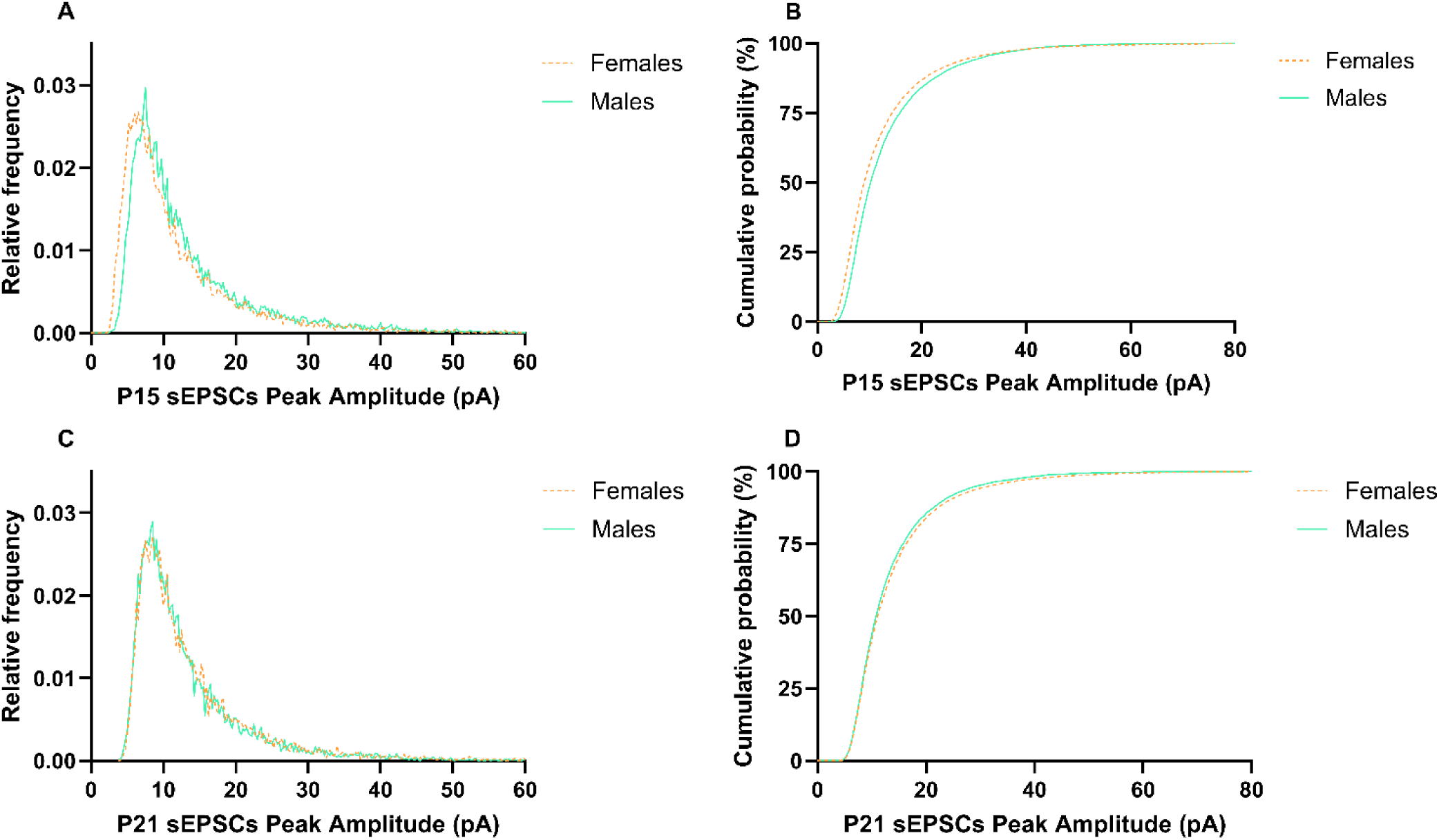
Distribution of sEPSC peak amplitudes at different early postnatal stages between sexes. Normalized histograms show the distributions of sEPSC peak amplitudes in females (orange/dashed) and males (green/solid) at P15 (**A**) and P21 (**C**). (**B, D**) Corresponding cumulative probability plots show the distributions of sEPSC peak amplitudes across sexes.

## Notes

### Competing Interest Statement

The authors have declared no competing interest.

### Summary of Updates

Added a link to the GitHub repository containing the Python script used to analyze the data.

